# Self-renewal of neuronal mitochondria through asymmetric division

**DOI:** 10.64898/2025.12.17.694973

**Authors:** Tejashree Pradip Waingankar, Camryn Zurita, Angelica E. Lang, Cliff Vuong, Ahmad Shami, Diana Bautista, Catherine Drerup, Samantha C. Lewis

## Abstract

Mitochondrial ATP production is essential for life. Mitochondrial function depends on the spatio-temporal coordination of nuclear and mitochondrial genome expression, yet how this coordination occurs in highly polarized cells such as neurons remains poorly understood. Using high-resolution imaging in mouse peripheral sensory neurons and zebrafish larvae, we identified a sub-population of mitochondria enriched in mtDNA that are positioned at the collateral branch points of long sensory neurites, both *in vitro* and *in vivo*. While the mitochondria in neurites are generally depleted of mtDNA, those at axon branch points preferentially engage in mtDNA replication and transcription, accumulate nuclear-encoded mitochondrial mRNA, and are spatially linked to nascent cytosolic peptide synthesis. The mtDNA-positive mitochondrial pool exhibits asymmetric genome partitioning at division, shedding highly motile daughters that lack mtDNA. Asymmetric division rejuvenates the membrane potential of the mtDNA-rich, biogenesis-dedicated mitochondria. We also found that, in peripheral sensory neurons, axonal mitochondria rarely fuse or share matrix contents, explaining how differentiated daughters maintain their distinct composition and fate after fission. Thus, division-coupled mitochondrial self-renewal is yoked to neurite topology in sensory neurons, patterning mitochondrial diversity and homeostasis from micron to meter scales.

## Main

Neurons are highly polarized and compartmentalized cells that require considerable amounts of energy, lipids, and amino acids for their growth and activity. In vertebrates, Dorsal root ganglia (DRG) extend the longest peripheral axons in the body, reaching up to 1.5 meters in humans to innervate peripheral tissues^1^. In vertebrates, primary afferent peripheral sensory neurons of the DRG are pseudounipolar and exclusively extend axons, which bifurcate into particularly long highly branching arbors^1^. This is in contrast to neurons in the central nervous system where soma typically extend neurite projections that may differentiate into either dendrites or axons.

The branch points of DRG neurites are important signaling hubs that mediate the afferent transmission of thermal, chemical, and mechanical information to the central nervous system, including signals perceived as itch or pain. Peripheral axons grow quickly in development and remodel their branching in response to intracellular signals as well as cues from the surrounding tissue they innervate. Mitochondria are key metabolic organelles that produce the ATP, lipids, neurotransmitter precursors, and Calcium buffering required for neurite growth^2–6^. As a result, the catabolic and anabolic functions of mitochondria are at the nexus of DRG homeostasis. A central question in neuronal biology is how mitochondrial integrity is maintained in neurites, where active transport of proteins and RNA from the soma is likely limiting for biogenesis.

Mitochondria harbor their own double-stranded DNA genome, which encodes essential subunits of the electron transport chain (ETC) and functional RNAs for their expression on mitoribosomes residing in the mitochondrial matrix^7^. Mitochondrial DNA (mtDNA) synthesis is a hallmark of mitochondrial biogenesis, coupled to mitochondrial membrane growth and division for distribution to daughter organelles^8,9^. Yet mitochondria rely exclusively on genes encoded in the nuclear genome for mtDNA synthesis and transcription, including all mitochondrial DNA and RNA polymerases, packaging, and transcription factors.

MtDNA is packaged into discrete protein-nucleic acid complexes termed mitochondrial nucleoids, which serve as the units of inheritance and transport of mtDNA within and between mitochondria^7,10^. Little is known about mitochondrial homeostasis in DRGs, relative to proliferating cells or CNS neurons. In proliferating cells, mtDNA synthesis, nucleoid partitioning, and mitochondrial fission are spatio-temporally coordinated to ensure mtDNA abundance scales with mitochondrial network volume^8,11^. Diffusion-limited mitochondrial nucleoids, marked by the essential mtDNA-packaging protein TFAM, are distributed throughout mitochondrial networks by fission^10,12^. A subset of nucleoids additionally recruits mtDNA polymerase Pol Gamma and/or RNA polymerase POLRMT, marking those mitochondria engaged in active biogenesis^10,12^. How mitochondrial integrity and mtDNA abundance are homeostatically maintained in post-mitotic neurons, especially in neurites hundreds or thousands of microns from the soma, is unclear.

In iPSC-derived human neurons, local translation of mitochondrial division receptor proteins is a means of regulating mtDNA-linked mitochondrial division in dendrites, likely in coordination with the local endoplasmic reticulum (ER)^13–16^. The long peripheral neurites of mammalian sensory neurons contain abundant mitochondria, ER, as well as the cytoplasmic machinery for local translation^17,18^. Moreover, messenger RNAs of the nuclear-encoded mitochondrial proteome are actively trafficked into axons on endo-lysosomes, and on the outer mitochondrial membrane itself, suggestive of in-neurite biogenesis programs^19–22^. These previous findings raise fundamental questions about where and when nuclear and mitochondrial biogenesis programs converge in neurites. However, where, whether, and when mtDNA synthesis occurs in neurites, especially the extremely long neurites from DRGs, has been elusive.

In this study, we use high-resolution microscopy to map the distribution, replication, expression, and trafficking of mitochondrial genomes between the soma and neurites of mouse and zebrafish peripheral sensory neurons, *in vitro* and *in vivo*, at the single-mitochondrion level. We identify a distinct mitochondrial pool positioned at neurite collateral branch points that is dedicated to mtDNA synthesis and expression, coincident with local RNA accumulation and cytoplasmic translation. We find that these biogenesis-devoted mitochondria are maintained by fission events characterized by asymmetric mtDNA partitioning; half of the differentiated daughter organelles lack mtDNA. Asymmetric division rejuvenates the biogenesis-dedicated mitochondrial pool, while daughters lacking mtDNA exhibit lesser membrane potential, and are rapidly trafficked from the site of biogenesis. These findings indicate that local biogenesis and fission maintain mitochondrial integrity far from the soma, thereby spawning genomeless mitochondria within neurites. We suggest a model in which mitochondrial diversity is integrated with neurite topology via cycles of mitobiogenesis and self-renewal to shape the bioenergetic landscape in peripheral neurites far from the soma.

## Results

To investigate how mitochondrial and mtDNA homeostasis are maintained in long neurites far from soma, we imaged mitochondria and mtDNA in primary murine DRG neurons marked by expression of tdTomato under control of the Pirt promoter^23–25^. We performed live laser-scanning fluorescence microscopy of tdTomato-positive DRG neurons co-stained for mitochondria (Mitotracker Deep Red) and DNA (SYBR Gold) and cultured them on glass-bottom dishes for 24 hours. We observed clear soma and bifurcated stem-axons distinctive to pseudounipolar neurons, which extended into periodically branching neurite arbors that were on average 750 microns long (21 neurons from 4 replicates; **Fig 1a; Extended Data FigS1a**). After initial stem-axon bifurcation, subsequent neurite branching occurred most frequently in a zone 40-100 microns from soma (**Extended Data FigS1b**). Abundant mitochondria were present throughout soma and neurites, some of which colocalized with SYBR Gold puncta that report mtDNA nucleoids. Mitochondria that were dense with mtDNA occupied the majority of the soma volume. In contrast, 49% of mitochondria within neurites lacked detectable mtDNA entirely, a difference that was apparent immediately after stem-axon bifurcation (**Fig. 1b-1c**).

**Figure 1:**
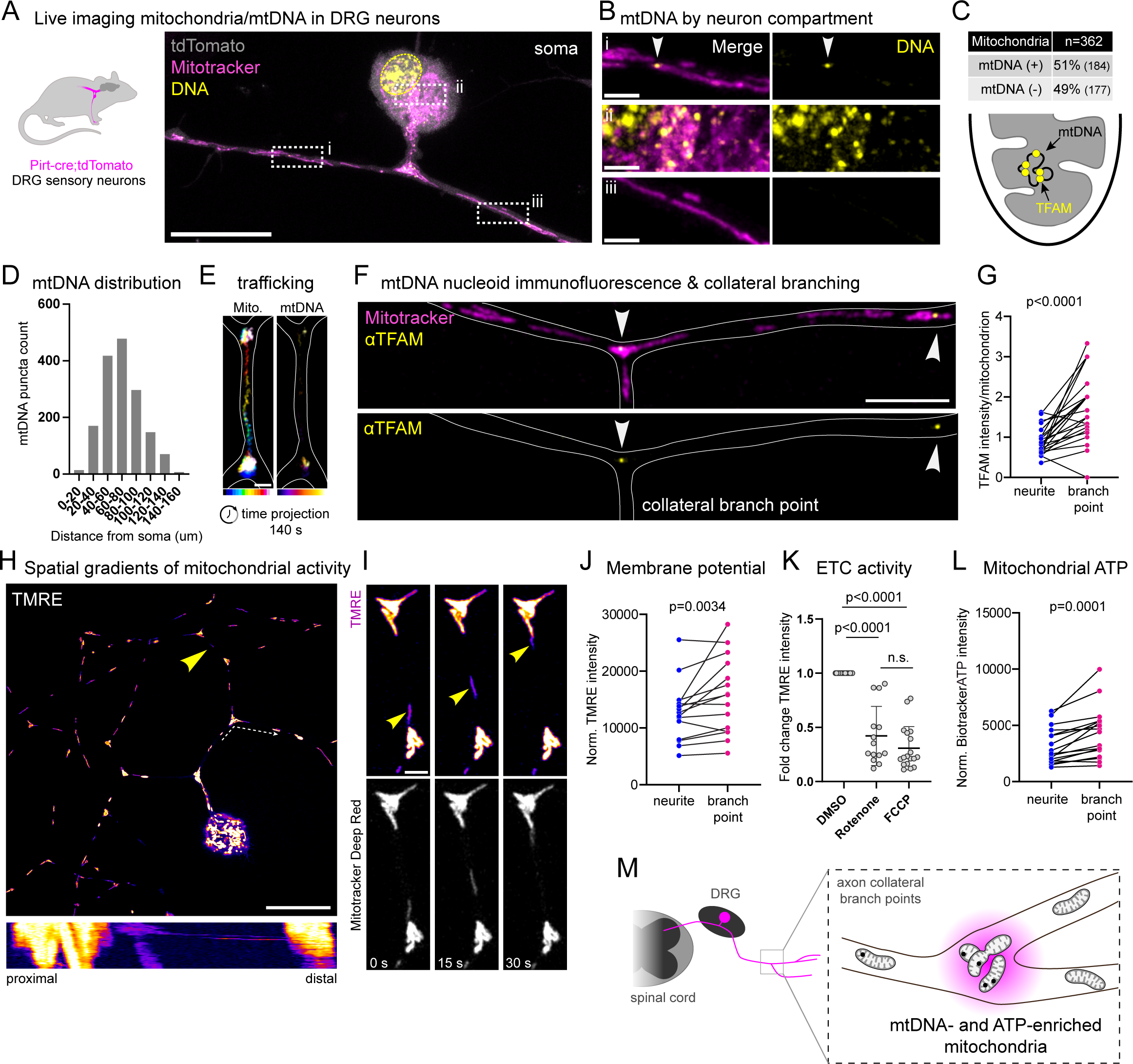
(A-B) Schematic illustrating the location of DRGs in the mouse model, along with the representative images of mitochondria (Mitotracker: magenta) and mtDNA (SYBR Gold: yellow) in neuronal sub-compartments. Scale bar: 20 and 2 µm. (C) Quantification of mitochondria with or without mtDNA within the neurite, excluding the soma. (D) Distribution of mtDNA along the length of the neurite in live DRG neurons labeled with MTDR and SYBR Gold, quantified relative to the cell body at the first timepoint of the time-lapse movie. (E) Color-coded projection through time of mitochondria and mtDNA trafficking. Scale bar: 2 µm. (F) Immunofluorescence image of sensory neurons with Mitotracker (magenta), EU (not shown), and anti-TFAM antibody (yellow). Scale bar: 5 µm (G) Quantification of TFAM puncta per mitochondrion. The total number of TFAM and mitochondria was quantified using Arivis. TFAM and mitochondria at branch points were manually counted and plotted to compare each field of view relative to the neurite. (H-I) Live cell imaging of neuronal mitochondria with Mitotracker (IMM: Inner Mitochondrial Membrane, white) and TMRE (Fire lookup table). Scale bars: 20 µm and 2 µm. (J) Mean TMRE intensity and corresponding area by Mitotracker were quantified at time points 1, 15, and 30, with equal numbers of neurite and branch-point mitochondria. The average intensity-to-area ratio at neurite and branch points is matched within the same field of view. (K) Live cell imaging of mitochondria (Mitotracker) and inner membrane potential (TMRE) of neurons treated with DMSO, Rotenone (30 µM for 3 h), and FCCP (20 µM for 1 h). TMRE intensities were quantified as mentioned in Fig. 1I. (L) Mean ATP intensity corresponding to MitoTracker green FM area was quantified at time points 1, 15, and 30 in an equal number of neurite and branch point mitochondria. The average ratio of ATP intensity to area for neurite and branch point mitochondria is plotted within the same field of view. (M) Schematic representation of heterogeneity at the branch point mitochondria.

We next quantified mitochondrial and mtDNA density along the initial 160 microns of neurites as they extended from the soma. The length of individual mitochondria was not correlated with distance from the soma centroid (**Extended DataFig. S1c**). In contrast, the density of mitochondrion-colocalized mtDNA rose steadily as the distance increased, peaking in a zone 40-80 microns from soma (**Fig. 1d**). Thus, mitochondria just distal to the stem-axon were relatively depleted of mtDNA, while mitochondria further from the soma were relatively enriched with mtDNA.

Peak mtDNA density coincided with the zone of peak neurite branching; thus, we asked whether mtDNA-rich mitochondria were spatially linked to the bifurcations of sensory afferents that form collateral neurites. Time-lapse microscopy revealed motile mitochondria trafficking anterograde and retrograde (**Fig 1e**). In contrast, mitochondria harboring mtDNA puncta were stably associated with neurite branch points (**Fig 1e; Supplementary Video 1; Extended Data Fig.S1d**). We next employed immunofluorescence microscopy to examine an orthogonal marker for mtDNA nucleoids, the essential mtDNA packaging and transcription factor TFAM, in Mitotracker-labeled mitochondria^12^. Discrete TFAM puncta were enriched within a subset of mitochondria, as previously reported for cortical neurons and oligodendrocytes^26,27^. Consistent with our direct visualization of mtDNA by SYBR Gold staining, mitochondria harboring TFAM puncta were proximal to neurite branch points (**Fig. 1f,g**). We again quantified the frequency of neurite branching by distance from soma, now paired with quantification of TFAM foci density in the same neurons, finding that neurite arborization and mitochondrial mtDNA density were indeed correlated (**Extended Data Fig.S1e).** These data indicate that mitochondria are differentially positioned along sensory afferents in accordance with their mtDNA content.

We wondered if mitochondrial bioenergetic activity may also vary by positioning along neurites. To examine spatial gradients of mitochondrial activity, we first co-labeled WT neurons with the cationic dye tetramethylrhodamine ethyl ester (TMRE) and the membrane potential-insensitive dye Mitotracker Deep Red as a control (**Fig. 1h**). Kymograph analysis revealed that TMRE fluorescence intensity varied between motile, presumably genomeless, mitochondria and adjacent stationary mitochondria (**Fig. 1i; Supplementary Video 2**). After normalization to Mitotracker Deep Red, mitochondria stationed at neurite branch points exhibited significantly higher TMRE intensity than other mitochondria in the same neurites (**Fig. 1j**). To test whether mitochondrial membrane potential is dependent on ETC function in sensory afferents, we compared these results to neurons treated with either the Complex I inhibitor rotenone or the ionophore Carbonyl cyanide-p-trifluoromethoxyphenylhydrazone (FCCP). All mitochondria exhibited reduced TMRE intensity upon rotenone or FCCP treatment, relative to vehicle (**Fig. 1k; Extended Data Fig.S2a**). Thus, in vitro, mitochondria in the sensory afferents of DRG neurons are dependent on Complex I function to maintain their membrane potential.

We next co-labeled WT neurons with Biotracker ATP, a switchable, small-molecule rhodamine derivative that exhibits a 5-fold increase in fluorescence upon reversible ATP binding and Mitotracker Green^28,29^ . Consistently, normalized Biotracker ATP fluorescence intensity was greater in the mitochondria localized to collateral branch points than those elsewhere in the same neurites (**Fig. 1l; Extended Data Fig.S2b**). Moreover, this difference remained stable over time (**Extended Data Fig.S2c**). Taken together, these data indicate that, while the neurite compartment is profoundly depleted of mtDNA, a pool of mtDNA-rich, high-activity mitochondria marks sites of sensory afferent collateralization (**Fig. 1m**).

Given that mitochondrial genome synthesis and expression are linked to expansion and quality control of the mitochondrial network, we considered whether mtDNA-enriched mitochondria specifically localized to neurite branch points represent a sub-population engaged in mitobiogenesis^8,10,11^. We pulse-labeled neurons with the click chemistry-compatible nucleoside analogs 5-ethynyl-2-deoxyuridine (EdU) or 5-ethynyluridine (EU) to label nascent DNA or RNA, respectively. We validated that EdU-labeled (**Fig. 2a**) and EU-labeled (**Fig. 2b**) puncta colocalized with TFAM and mitochondria at neurite branch points^30^ (**Extended Data Fig.S3a,b**). The subset of mtDNA nucleoids engaged in synthesis or transcription was significantly enriched in the mitochondria stationed at neurite branch points (**Fig. 2c,f).** Both EdU and EU signals were sensitive to inhibition of the sole mitochondrial RNA polymerase, POLRMT, which both primes mtDNA replication and executes processive polycistronic transcription and served as a control (**Fig. 2d,g**). To assess the proportion of mtDNA nucleoids engaged in synthesis or transcription at neurite branch points, we compared the number of labeled mtDNA nucleoids per TFAM focus (**Fig. 2e,h; Extended Data Fig.S3a,b**). Consistent with our other data, the pool of mitochondria at branch points exhibited a greater number of TFAM puncta, of which a greater proportion were labeled by EdU or EU. These assays provide evidence that not only are mitochondria at neurite branch points enriched with mtDNA, but are also preferentially engaged in mtDNA synthesis and/or transcription.

**Figure 2:**
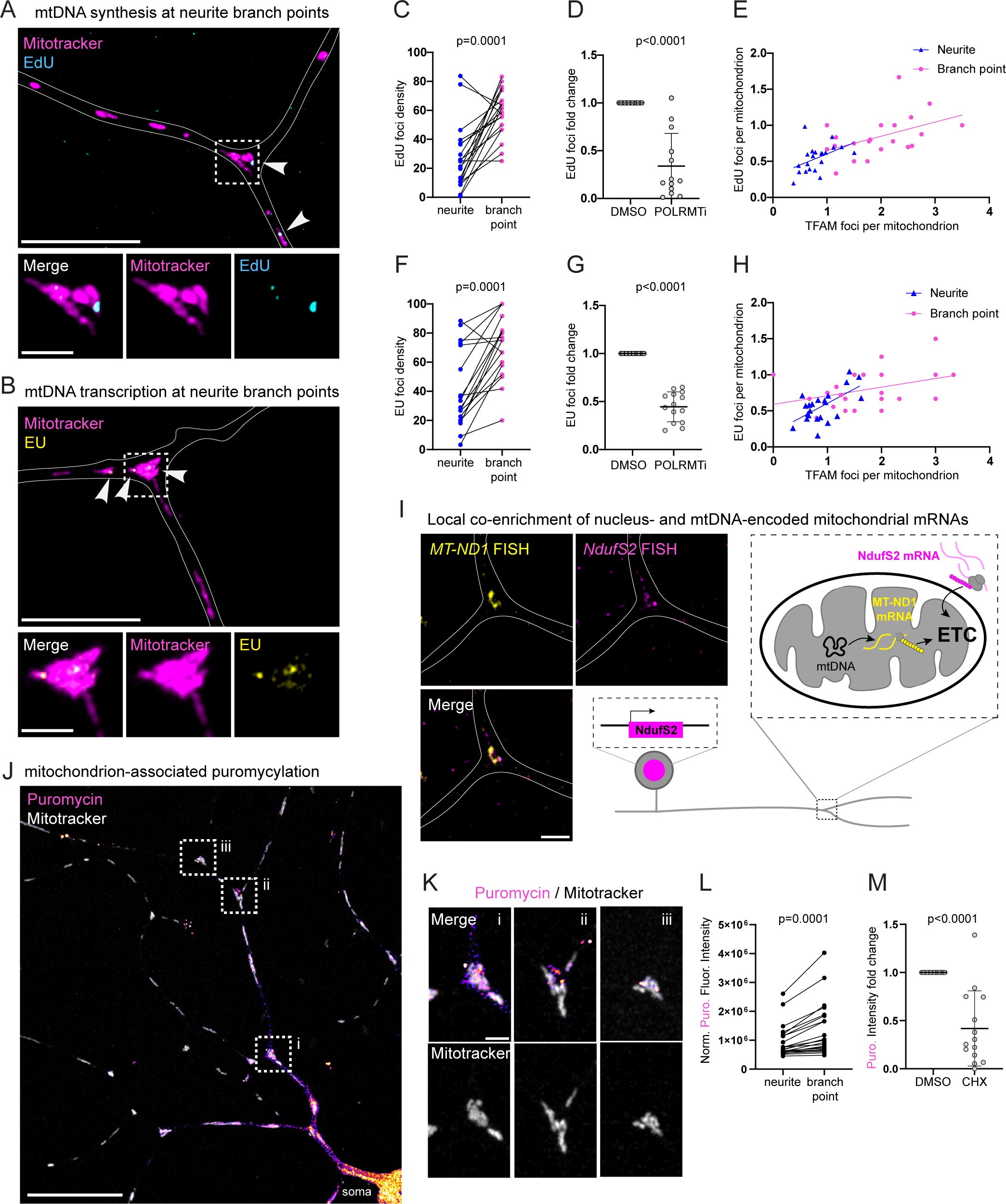
(A) Metabolic labeling of mtDNA replication, mitochondria (Mitotracker: magenta), EdU (blue). Scale bar: 20 and 2 µm. (B) Metabolic labeling of mitochondrial transcription, mitochondria (Mitotracker: magenta), EU (yellow). Scale bar: 20 and 2 µm. (C and F) Quantification of EdU (C) and EU (F) foci per mitochondria. The total mitochondrial and EdU/EU counts were quantified using Arivis. Branch point mitochondria were manually assessed for the presence of EdU/EU foci. The proportion of mitochondria at the branch points or neurites that contained EdU/EU per field of view is plotted. (D and G) Quantification of immunofluorescence analysis of neurons treated with DMSO or POLRMTi (IMT1B: 10 µM for 96 h) followed by Mitotracker and EdU (D) or EU (G) labeling. (E) Quantification of EdU per mitochondria with respect to TFAM per mitochondria at the neurite and branch points. R square for neurite: 0.2029, R square for branch points: 0.2166. (H) Quantification of EU distribution with respect to TFAM in the neurite and branch point mitochondria. R square for neurite: 0.3837, R square for branch point: 0.1413. (I) RNA-FISH analysis of neurons labeled with Tom20 (not shown), MT-ND1 (yellow), and Ndufs2 (magenta). Scale bar: 2 µm. Schematic illustrating the assembly of ETC complexes away from the cell body. (J and K) Immunofluorescence image of neurons labeled with Mitotracker (gray) and anti-puromycin antibody (fire lookup table). Scale bar 20 and 2 µm. (L) Quantification of the sum puromycin intensity normalized to the area, compared between neurites and branch points within the same field of view. (M) Quantification of the sum puromycin intensity normalized to the area for DMSO and cycloheximide (50 µg/ml for 1 h).

Approximately 99% of the mitochondrial proteome is encoded in the nuclear genome, thus mitochondrial mRNAs are actively trafficked from soma into neurites for local translation across a variety of neuronal types^31^. Given our observation of active mtDNA transcription, we wondered whether mitochondrial mRNA encoded in the nucleus could be targeted to neurite branch points for co-expression. We directly visualized the spatial distribution of two mitochondrial mRNAs required for complete assembly of Complex I by RNA-FISH, one encoded in the nuclear genome, and one encoded in mtDNA (**Fig. 2i; Extended Data Fig.4a**). We selected *Ndufs2* and *MT-ND1* for these experiments, as mutations in either gene cause axonal neuropathy and Parkinsonism in mouse models, indicating that their regulation is essential for neurite maintenance^32,33^. *NdufS2* and *MT-ND1* FISH signals colocalized with each other (**Fig. 2i**) and with the mitochondrial outer membrane marker TOM20 at neurite branch points (**Extended Data Fig.S4b**). We quantified the normalized density of RNA-FISH signals in neurite mitochondria, finding that those organelles positioned at neurite branch points were significantly enriched (**Extended Data Fig.S4c**).

Given these findings, we speculated that local cytosolic translation in neurites could mark the coordinated expression of mitochondrial genes from both genomes. We examined whether nascent peptides also accumulated at neurite branch points using puromycylation labeling^34^. Anti-puromycin immunofluorescence in Mitotracker-labeled neurons revealed that soma were highly enriched for labeled peptides, relative to neurite labeling (**Fig. 2j**). However, we noticed hotspots of labeled nascent peptides that colocalized with mitochondria, including at the branch points (**Fig. 2k,l; Extended Data Fig.S4d**). The fluorescence intensity of these punctate anti-puromycin signals was significantly reduced by inhibiting cytosolic translation with cycloheximide, validating that the signal correlated with active translation (**Fig. 2m**). Altogether, these data show that mitochondrial genome synthesis and transcription coincide with local accumulation of nucleus-encoded mitochondrial mRNA and nascent translation products at neurite branch points. This suggests that these mitochondrial hubs could preferentially function in Complex I biogenesis.

To determine whether a distinct, mtDNA-rich mitochondrial population could mark neurite branch points *in vivo*, we co-imaged mitochondria and mtDNA nucleoids in axons of live zebrafish sensory neurons. We specifically examined the transport and positioning of TFAM (as a general nucleoid marker), or POLG2 (a subunit of the mitochondrial DNA polymerase and marker for mtDNA synthesis) in pLL lateral line neurons. These pLL neurons contain abundant mitochondria within a long, periodically branching peripheral axon, are found just under the skin, and are mature in transparent larvae at 4 days post-fertilization (dpf), and thus highly amenable to live imaging^35–37^.

We injected DNA plasmids encoding two genes separated by a degradable 2A linker into zygotes for mosaic expression, one a mitochondrial matrix reporter, and the other encoding either TFAM-mRFP or POLG2-GFP (**Fig. 3a**). Similar to observations in primary mouse DRG neurons, TFAM-mRFP fluorescence localized to axonal branch points *in vivo*, significantly more so than in other mitochondria (**Fig. 3b,c**). POLG2 puncta were also significantly enriched amongst the population of mitochondria at branch points, and to an even greater extent (**Fig. 3d**). We next leveraged this system to ask how stable the localization of POLG2-GFP puncta to axon branch points was in larvae. In these experiments, we co-imaged mitochondria and POLG2-GFP once per minute for 5.5 hours (**Fig 3.e; Extended Data Video 3**). Both anterograde- and retrograde-moving mitochondria trafficked through axon branch points; the majority of which lacked POLG2-GFP puncta. In contrast, a subset of mitochondria that harbored prominent POLG2-GFP signals remained continuously anchored at each site of branching for multiple hours, even as the axon and most proximal synapse were actively remodeled (**Extended Data Video 3; Fig.3f**). These results indicate that mtDNA nucleoid enrichment and synthesis at axon branch points is a conserved feature of sensory neurons both *in vitro* and *in vivo*.

**Figure 3:**
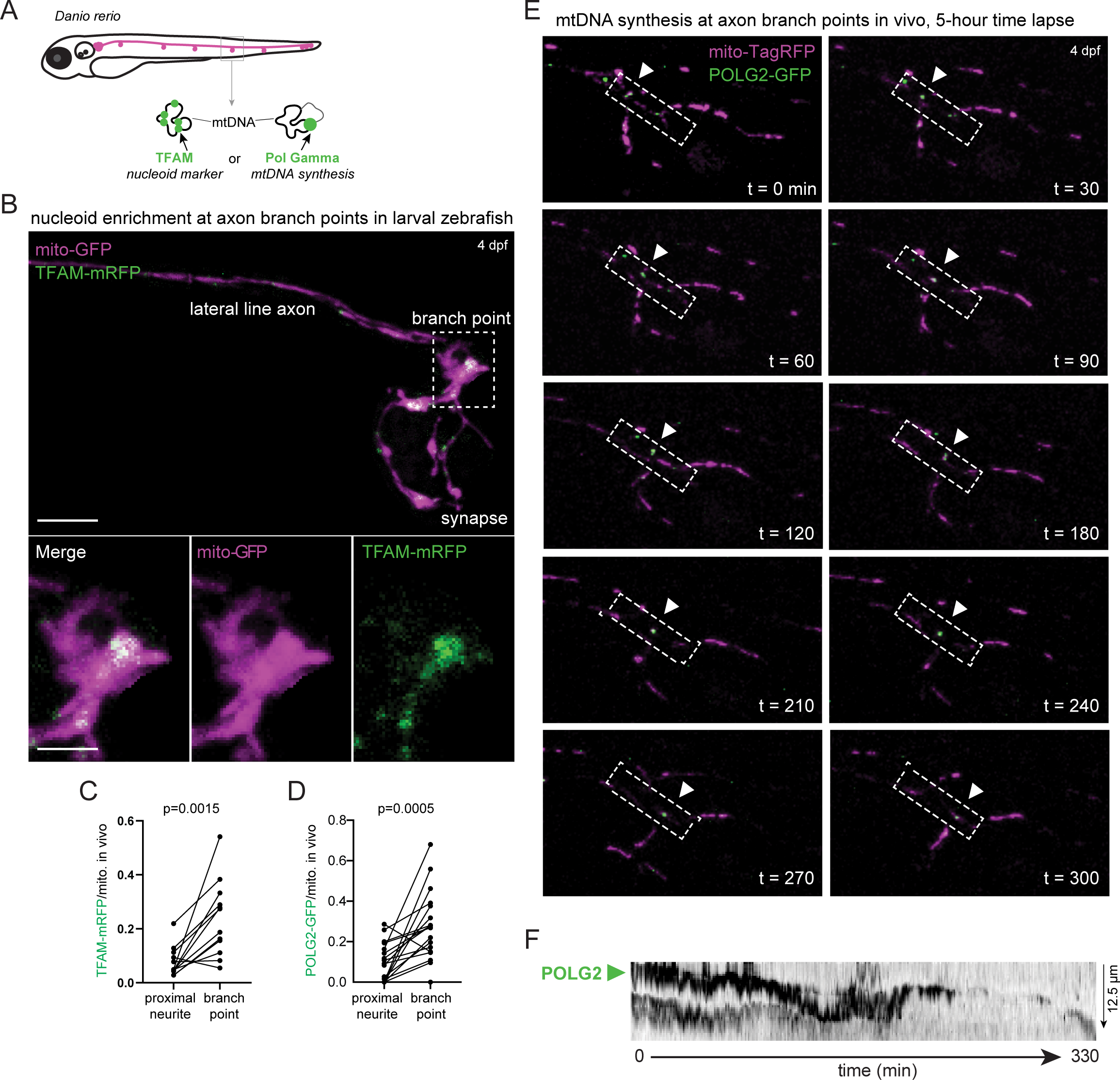
(A) Schematic illustration of zebra fish expressing markers for mitochondria, TFAM, and POLG2. (B) live cell imaging of zebra fish expressing TFAM-mRFP (green) and neurod:mito-GFP (magenta). Scale bar 5 µm. (C) Quantification of TFAM-mRFP volume to mitochondrial volume in zebra fish neurons. (J) Quantification of POLG2-GFP volume to mitochondrial volume in zebra fish neurons. (E) Live cell imaging of zebra fish expressing POLG2-GFP (green) and neurod:mito-RFP (magenta). Representative time points of a pLL axon terminal expressing 5kbneurod1: Polg2-GFP-p2a-mitoTagRFP. The boxed region indicates the region used for the kymograph. Arrowhead indicates puncta at the branch point. (F) Kymograph of POLG2 movement in the boxed regions of Figure 3E.

We next investigated the fusion, fission, and motility dynamics of mitochondria in neurites, using mouse DRG neurons in which the photoconvertible green fluorescent protein Dendra2 was targeted to the mitochondrial matrix under control of the Pirt promoter^38^ **(Fig. 4a**). Our observation that the pool of mitochondria anchored at branch points maintains their distinct protein and nucleic acid composition over time led us to speculate that they could exhibit distinct mitochondrial dynamics as well. Upon gentle photoconversion of mitochondria with UV-violet light, cox8a-Dendra2 fluorescence is rapidly converted from green to red. We calculated the red: green fluorescence ratio after photoconversion to be 40-50%, depending on the experiment, which was stable over the subsequent minutes of time-lapse imaging (**Fig. 4a,b**).

**Figure 4:**
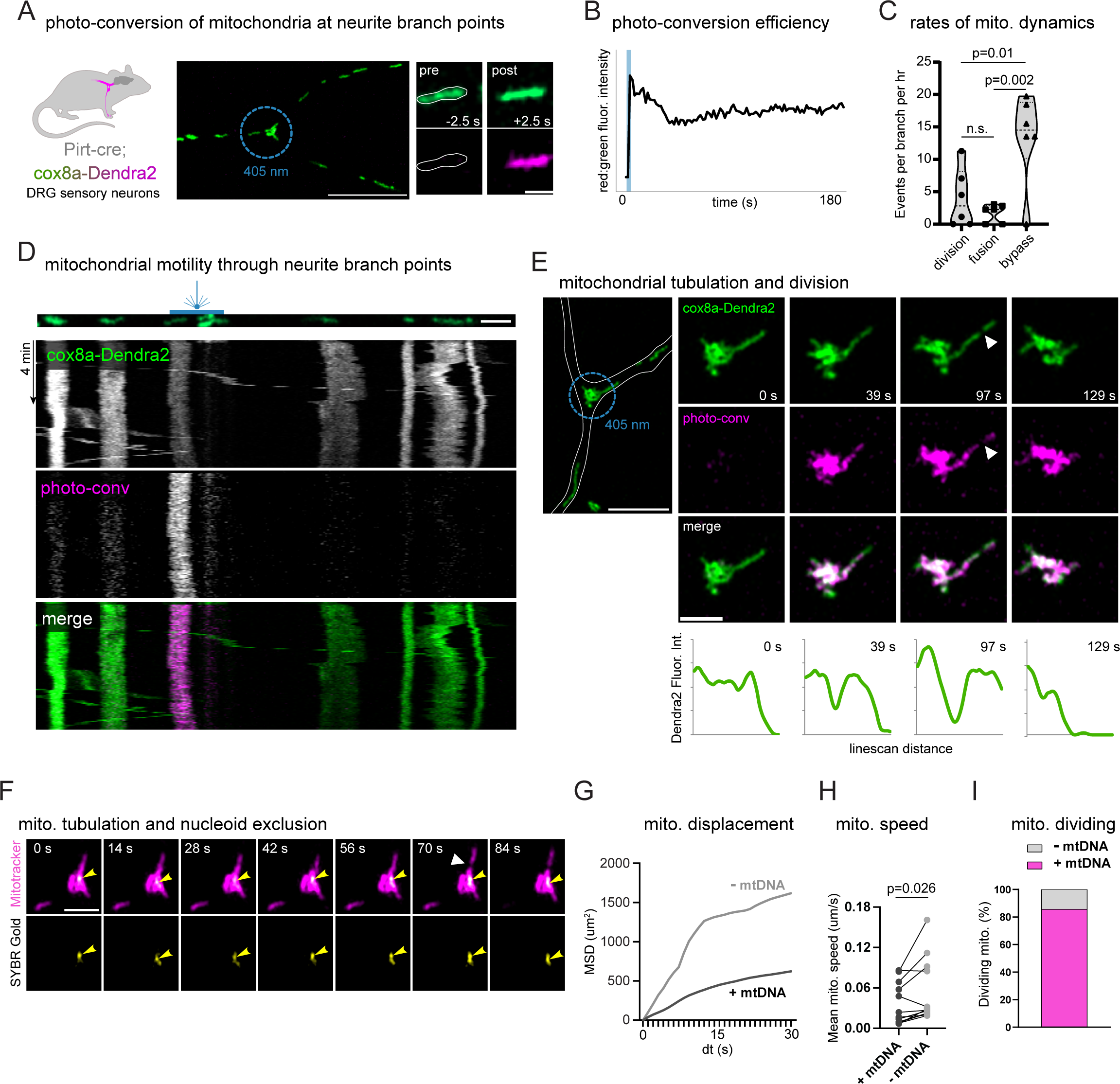
(A) Representative images of Cox8a-Dendra2 photoconversion at the branch points. (B) Quantification of red-to-green fluorescence intensity post-conversion over time. (C) Quantification of division, fusion, and bypass events at the branch points over the time course of one hour (fusion: neurite mitochondrion enters the branch point and stays over the course of the movie, division: branch point mitochondria divide to send out one of the daughters in the neurite, bypass: neurite mitochondria enter the branch point and leaves without exchanging the contents with the branch point mitochondria). (D) Kymographs of the branch-point mitochondria, pre-(green) and post-conversion (magenta), to visualize fusion and bypass events. (E) Live-cell imaging of mitochondria at branch points, pre-(Cox8a-Dendra2: green) and post-conversion (magenta), to show a representative division event. Scale bar: 2 µm. Line-scan analysis at branch points to represent Dendra2 intensity before and after division over time points. (F) Live cell imaging of mitotracker (magenta) and SYBR Gold (yellow) to visualize asymmetric division events at the branch points. (G) Cumulative mean squared displacement was calculated from tracks of mitochondria with or without nucleoids (183 and 182 tracks, respectively) in live DRG neurons labeled with MTDR and SYBR Gold for at least 10 time points. (H) Max track speed was measured from tracks of mitochondria with or without nucleoids (183 and 182 tracks, respectively) in live DRG neurons labeled with Mitotracker Deep Red and SYBR Gold for at least 10 time points. The maximum track speed was then averaged across neurons. (I) Quantification of the number of divisions in the neuronal mitochondria with or without mtDNA.

We used this cox8a-Dendra2 diffusion assay to determine the frequency of matrix content mixing upon fusion or separation upon fission, by imaging every 2.5 seconds for 4-10 minutes after photoconversion (**Fig. 4c; Supplemental Video 4**). We readily observed anterograde and retrograde mitochondrial trafficking. Yet, detectable fission and fusion were rare events, regardless of whether mitochondria were localized to branch points or elsewhere in neurites (**Extended Data Fig.S5a**). From cumulative analyses of experiments in 43 neurons (6 animals), we estimated the frequency of mitochondrial division or fusion to be 0-3 events per neurite branch, per hour (**Fig. 4c**). We observed that 10-15 mitochondria trafficked through each branch point in the same timeframe, indicating that while mitochondria had opportunities for content mixing, they typically abstained from doing so (**Fig. 4c**). We visualized the stability of mitochondrial positioning in neurites using kymograph analysis, confirming that most actively trafficking mitochondria do not appreciably interact with photoconverted organelles (**Fig. 4d**).

Interestingly, we found that when mitochondrial division did occur, it was preceded by the stereotyped extension of a mitochondrial tubule from the anchored mitochondria at the branch point. These tubules underwent subsequent constriction and division, after which a small daughter mitochondrion rapidly trafficked away from the stationary pool (**Fig. 4e**). We observed similar events in neurons labeled with Mitotracker and SYBR Gold (**Fig. 4f**). Strikingly, we found that in 91% of these division events, the motile daughter mitochondrion lacked mtDNA (**Extended Data Fig.5b**). This prompted us to quantify the mean squared displacement (MSD) of individual mitochondria with or without mtDNA in neurites, finding that genomeless mitochondria are significantly more motile than those with at least one mtDNA nucleoid (**Fig. 4g**). Genomeless mitochondria exhibited greater speed of motility in neurites, which likely accounts for this difference (**Fig. 4h**). These data indicate that the mitochondrial pool anchored at neurite branch points largely abstains from matrix content mixing with motile mitochondria that pass by. When fission does happen, mtDNA partitioning is asymmetric, generating highly motile, genomeless daughters.

We wondered if the genomeless mitochondria generated via asymmetric division could be a source of the highly motile yet low membrane potential mitochondria we had previously observed (**Fig. 1**). We hypothesized that the biogenesis-dedicated mitochondrial pool at neurite branch points could periodically shed low-quality fragments, which could serve some other function and/or be degraded elsewhere in the neuron. Consistent with this idea, we noted that genomeless mitochondria divided much less frequently than those containing at least one mtDNA nucleoid (**Fig. 4i**).

To address this, we imaged mitochondria in murine DRG neurons co-labeled by TMRE and Mitotracker Deep Red once more, this time focused on capturing the rare mitochondrial division events at neurite branch points (**Fig. 5a**). Consistent with our previous experiments, extension of a distinct mitochondrial tubule preceded constriction and division, generating a small mitochondrial fragment (**Fig. 5b**). Pre-division tubules contained some inner mitochondrial membrane, as they were labeled by both dyes, yet exhibited significantly lower membrane potential as reported by TMRE fluorescence intensity, even normalized for their small size (**Fig. 5c**). Here, we represent these data as a comparison of the normalized TMRE values for the mitochondrial mass pre-division (“mother”), to each of the masses generated (“daughters” that either remain or exit the branching site). This shows that normalized TMRE intensity in “mother” mitochondria increases upon fission, suggesting that they are rejuvenated in this process. We repeated this experiment, but in neurons co-labeled with BiotrackerATP and Mitotracker Green, with similar results (**Fig. 5d,e,f**). Stationary mitochondria at neurite branch points shed small daughters characterized by low BiotrackerATP fluorescence intensity, while the remaining mitochondrial mass maintained or increased in this intensity.

**Figure 5:**
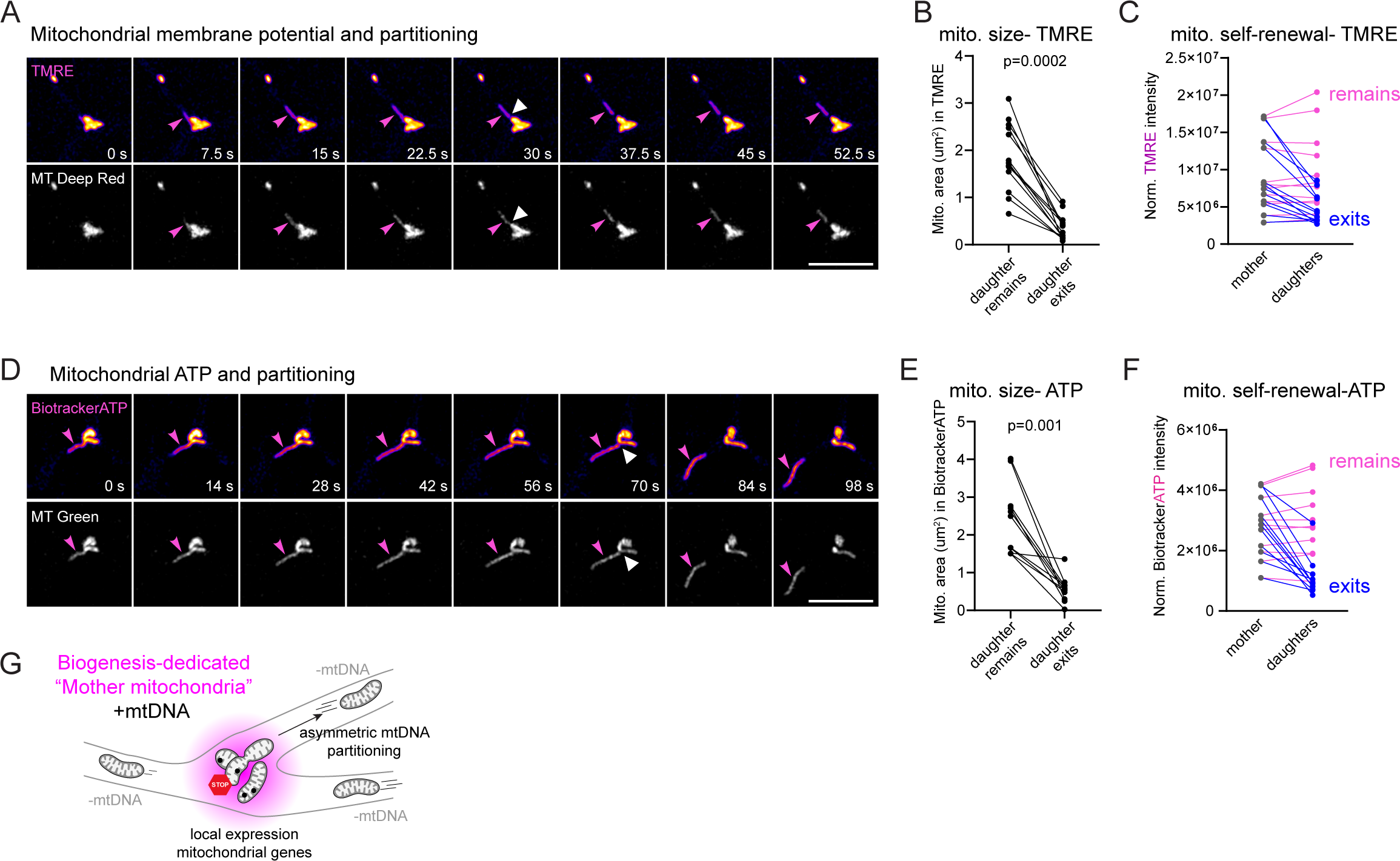
(A) Live cell imaging of TMRE (fire lookup table) and MitoTracker DeepRed (MT Deep Red: gray) to visualize membrane potential during divisions at the branch points. Scale bar: 5 µm. (B and C) Quantification of TMRE intensity (C) normalized to the mitochondria area (B) quantified from the MT Deep Red channel. The normalized intensity of the branch point mitochondria before division (mother) is compared with that of the daughters after division (remains: stays at the branch point after division, exits: leaves the branch point and enters a neurite). (D) Live cell imaging of BioTracker ATP (fire lookup table) and MitoTracker Green (MT Green: grey) to visualize divisions at the branch points. Scale bar: 5 µm. (E and F) Quantification of the BioTracker ATP intensity (F) normalized to the mitochondrial area calculated from the MT Green channel (E). The normalized intensities during divisions are plotted in the graph (mother: before division, remains: stays at the branch point after division, exits: leaves the branch point and enters a neurite).

We conclude that mitochondria anchored at neurite branch points comprise a biogenesis-active mother-mitochondrion pool that periodically differentiates lower quality, genomeless daughters through asymmetric fission (**Fig. 5g**). The distribution and motility of low-quality motile daughters is striking and may explain the stereotyped spatial patterning of mitochondrial diversity in DRG neurites. For example, cycles of asymmetric fission may facilitate the self-renewal of highly bioenergetically active mitochondria far from the soma, while generating a functionally distinct, genomeless class in the neurite.

## Discussion

Our study demonstrates a previously unappreciated organization of mitochondrial populations within peripheral sensory neurons, revealing how mitochondrial composition, dynamics, and genome content are spatially coordinated with local neurite topology and function. Mitochondrial dysfunction is broadly linked to neurodegenerative and neuromuscular disease, underscoring the importance of understanding how mitochondrial diversity and quality are regulated in neurons^39–41^. Our findings provide insight into how mitochondria are locally specialized within highly polarized neurite arbors.

It has been known for many years that DRG neurons contain multiple co-resident mitochondrial populations that differ in size and apparent density by electron microscopy^42^. We identified a high-activity, mtDNA-rich mitochondrial pool that specifically marks sites of sensory afferent collateralization. Not only are mitochondria at neurite branch points enriched for mtDNA, but this subset of mtDNA nucleoids is preferentially engaged in synthesis and/or transcription suggesting that the licensing of mtDNA maintenance processes within mitochondria is itself spatially regulated. Importantly, mtDNA nucleoid enrichment and active synthesis at axon branch points are conserved features of sensory neurons observed both *in vitro* and *in vivo*.

Our results further demonstrate that the mitochondrial pool anchored at neurite branch points largely abstains from matrix content mixing with the motile mitochondrial population. When fission events do occur within this anchored pool, mtDNA partitioning is asymmetric, producing highly motile daughter mitochondria that lack detectable mtDNA. These observations support a model in which mitochondria anchored at neurite branch points comprise a biogenesis-active “mother” pool that periodically differentiates lower-quality, genomeless daughters through asymmetric fission. Our findings are consistent with emerging evidence that genomeless mitochondria are likely depleted of the typical mitochondrial proteome. Across multiple neuronal types, axonal mitochondria are generally depleted of mtDNA and of proteins involved in mtDNA expression and broader electron transport chain assembly^30,43^. There may be immunological and physiological advantages to limiting the size of the biogenesis-dedicated mitochondrial pool within neurites to the minimal mass that is required to sustain compartmental function. For example, restricting neurite oxidative phosphorylation has been invoked as a mechanism to tune neuronal excitability in vivo by limiting reactive oxygen species generation^44,45^. Similarly, limiting mitochondrial biogenesis and, consequently, mtDNA content in neurites may reduce neuronal susceptibility to inflammation driven by the cytosolic release of mitochondrial nucleic acids to the cytoplasm. Mitochondrial DNA and RNA are recognized as foreign in the cytosol, activating RIG-I and cGAS-STING innate immune pathways likely to drive neuroinflammation in aging, making their excess a potential liability^46–48^.

Despite this apparent mtDNA depletion, mitochondrial genome expression and OXPHOS remain critical for neuronal health, as deficiencies in mitochondrial gene function (whether encoded in the nucleus or mtDNA) lead to axon degeneration, neuropathy, and Parkinsonism^49^. Our results demonstrate that mitochondrial diversification in composition and function tunes mitochondrial pools to local demands within neurites. We suggest that, in peripheral sensory neurons, the bioenergetic and functional landscape of neurites is shaped by mitochondrial diversity in accordance with branching topology, thereby integrating cellular function across length scales ranging from microns to meters.

The mtDNA-containing mitochondrial pool may also function to support local protein synthesis. Local translation is spatially linked to mitochondria and to axon branching propensity, and previous work has shown that translational machinery and respiring mitochondria are recruited to presumptive branch sites^50,51^. Axon branching is energetically demanding, requiring ATP to drive cytoskeletal remodeling, membrane expansion, and localized translation necessary to shape the secretory proteome^52,53^. Our results suggest that a key distinction between mitochondria that traffic past future branch sites and those that are selectively recruited may lie in their mtDNA content. Consistent with this idea, we observe that both mitochondrial mRNAs and nascent peptides accumulate in close proximity to mtDNA-positive mitochondria at neurite branch points.

These findings raise the question of how matrix-localized mtDNA content might be recognized or communicated to the exterior of the organelle. Greater motility of mitochondria lacking mtDNA has been previously observed in *Saccharomyces cerevisiae*^54,55^ and in *Arabidopsis*^56^. In fungi, it has been proposed that mtDNA nucleoids are connected to motor complexes or the cytoskeleton via the ERMES complex, which tethers ER and mitochondrial membranes at stable membrane contact sites^55,57^. However, the components of the ERMES complex are not conserved in vertebrates, leaving the identity of a functionally analogous complex unresolved in murine neuronal systems. Local actin patches have been implicated in axon branch site selection and also play roles in regulating the structure and dynamics of ER–mitochondria contacts where mtDNA synthesis occurs^8,58,59^. Proteins localized at ER–mitochondria contacts could further link the spatio-temporal specificity of mitochondrial activity to mitochondrial division and calcium release, providing a potential framework for coordinating mtDNA replication, fission, and functional output^60–62^. These hypotheses remain to be tested.

In summary, our findings demonstrate that asymmetric mitochondrial division facilitates the self-renewal of highly bioenergetically active mitochondria far from the soma, while simultaneously generating a functionally distinct, genomeless mitochondrial class within neurites. Our results suggest that the subset of mtDNA-enriched mitochondria are preferentially recruited to sites of neurite branching. This connection has broad implications for our understanding of the cellular pathology underlying human diseases, most obviously in the context of axonal degeneration and peripheral neuropathy, which are linked to defects in the same mitochondrial mRNAs we have mapped to “mother mitochondria” in this study. Our data suggest that pathogenesis in neuropathy, and potentially many more human diseases linked to mitochondrial dysfunction, could result primarily from disruption of the size or integrity of the “mother mitochondria” pool. Testing this model may provide a conceptual framework for understanding how mitochondrial diversity is generated and maintained in highly polarized neurons to support local function while preserving global cellular homeostasis, and how to recover mitochondrial integrity once lost.

## Online Methods

Manufacturer information for reagents, catalog numbers, and experimental details are listed in the **Extended Data Tables 1-5**.

### Animals and DRG culture

All mice were housed under standard conditions, as approved by the Animal Care and Use Committee of the University of California, Berkeley (14-hour light: 10-hour dark cycle, 21°C). C57BL/6J mice were obtained from The Jackson Laboratory. All experiments were performed under the policies and recommendations of the International Association for the Study of Pain and approved by the University of California, Berkeley Animal Care and Use Committee. The mouse line, Pirt-Cre (Pirttm3.1(cre)Xzd), used in the study to make photoconvertible GFP animals, was a gift from Dr. Xinzhong Dong (Johns Hopkins University, Baltimore, MD).

Additionally, PhAM floxed (B6;129S-Gt(ROSA)26Sortm1(CAG-COX8A/Dendra2)Dcc/J) animals were purchased from The Jackson Laboratory.

All experiments were performed on mice between 4 and 8 weeks old. The DRGs were isolated and cultured according to a previously published protocol^63,64^. Briefly, DRGs dissected from mice were incubated with prewarmed HBSS without Ca^2+^ and Mg^2+^ (Thermo Scientific), containing 2 mg/ml Collagenase P (Sigma-Aldrich) and 1 mg/ml Dispase (Sigma-Aldrich) for 45 min at room temperature (RT). The culture was centrifuged at 1000 rpm for 5 min at RT. The DRGs were treated with prewarmed 1 ml 0.25% Trypsin (Gibco) with agitation for 90 sec, followed by immediate Trypsin neutralization with 1 ml of complete MEM (minimal essential medium (Gibco), 10% horse serum (ThermoFisher Scientific), 1% penicillin/ streptomycin (Gibco), 1% L-Glutamine (Gibco), 1% MEM vitamin supplement (HyClone). The pellet was triturated 3 times with a fire-polished glass Pasteur pipette in a complete MEM containing DNase I (Thermo Scientific). Cells were spotted on 35 mm imaging dishes (Mattek) pre-coated with Laminin (1:100 in PBS). Finally, the cells were flooded with complete MEM after 2 h and incubated at 37□, 5% CO_2_ until the experiment.

### Microscopy, live cell imaging, and immunofluorescence

All the images and movies were acquired using a Zeiss LSM 980 with an Airyscan two-laser scanning confocal microscope. The microscope features lasers 405, 488, 561, and 639 nm, as well as a fast Airyscan detector array. Images were captured using an inverted 63X oil objective.

Live cell imaging was performed in a humidified chamber at 37°C and 5% CO_2_. Finally, Zeiss ZEN Blue software version 3.7 (Carl Zeiss) was used for Airyscan processing and maximum intensity projection.

For live-cell imaging, cells were cultured for 20-24 h, then incubated with the imaging dyes at 37□ and 5% CO_2_. The concentration and time of incubation for the dyes are as follows: MitoTracker Deep Red (ThermoFisher Scientific, 50 nM for 20 min), SYBR Gold nucleic acid stain (ThermoFisher Scientific, 1X for 20 min), Tetramethyl rhodamine, Ethyl Ester (TMRE, ThermoFisher Scientific, 200 nM for 20 min), BioTracker ATP Red (Sigma-Aldrich, 10 µM for 30 min), MitoTracker Green FM (ThermoFisher Scientific, 50 nM for 20 min).

Neurons were cultured for 20-24 h unless mentioned otherwise before being fixed for immunofluorescence. Cells were fixed in pre-warmed 4% paraformaldehyde (Electron Microscopy Sciences) for 20 min at RT, protected from light. The reaction was stopped by adding 0.2 M glycine for 5 min at RT. Cells were then permeabilized with 0.1% Triton X-100 in DPBS for 10 min, followed by washing with the blocking buffer, 0.1% TBST (1X TBS pH 8, 0.1% Tween 20) containing 1% BSA. The primary antibodies were added at the concentrations mentioned in the supplementary information and incubated overnight at 4□ for TFAM (Abcam) and Complex I (Invitrogen), and for one hour at RT for Tom20 (Proteintech). Samples were washed once with 0.1% TBST, then incubated with the secondary antibody at 1:2000 for 1 h at RT, washed with a blocking buffer, and imaged in DPBS. The primary and secondary antibodies used are listed in the supplementary information.

### EdU and EU labeling

Metabolic labeling of mitochondrial replication and transcription was performed using the manufacturer’s Click-IT EdU protocol with some modifications (Thermo Fisher Scientific).

For EdU labeling, cells were metabolically labeled with a 10 µM pulse of EdU for 3 h, followed by a 5 min chase in culture medium.

For EU labeling, cells were incubated with 1 µM triptolide for 1 h, followed by a 3 h pulse with 500 µM EU and a 5 min chase in culture medium.

For both EdU and EU, after 2 h pulse, mitochondria were stained with MTDR (50 nM for 1 h at 37□) before fixation with prewarmed 4% paraformaldehyde for 20 min at RT. The reaction was stopped by adding 0.2 M Glycine for 5 min at RT. Cells were permeabilized with 0.1% Triton X-100 in DPBS for 10 min, followed by 5 min incubation with blocking buffer. The click reaction with Alexa Fluor 488 picolyl azide was set up according to the manufacturer’s instructions for 30 min at RT. Cells were washed with a blocking buffer for 10 min at RT and imaged in DPBS.

For the IMT1B control, cells were treated with either vehicle control or IMT1B for 96 h at 37 □ then labeled with EdU or EU.

#### RNA FISH

RNA FISH was performed according to the previously published protocol with minor modifications^65^ (Biosearch technologies). All the probe sequences are available in the supplementary information. Briefly, for mt-RNR2, cells were incubated with 50 nM MTDR for 1 h before fixation. For Complex I FISH, an anti-Tom20 antibody (1:1000 for 1 h at RT) was used to visualize mitochondria. RNA-FISH probes were incubated for 16 h, then fixed with pre-warmed 4% paraformaldehyde. Other steps were performed as reported in^65^.

### Puromycin assay

Active cytoplasmic translation was analyzed by using puromycin as per the previously published protocol^13^. Briefly, after 20-24 h in culture, neuronal mitochondria were stained with MTDR (50 nM for 1 h) at 37 □, followed by incubation with 2 µM puromycin (Sigma-Aldrich) for 10 min.

Cells were washed once with culture medium and then fixed according to the protocol mentioned above. Neurons were incubated with anti-puromycin Alexa Fluor conjugate 488 for 16 h at 4 □. Cells were washed with a blocking buffer and imaged in DPBS.

For the negative control, cells were incubated with 50 µg/ml cycloheximide for 1 h before puromycin addition.

### Photoconversion of live neurons

DRGs expressing mitochondria-localized photoconvertible GFP were subjected to bleaching at 405 nm, and time-lapse movies were acquired at 488 nm and 561 nm. For bleaching in the region of interest (ROI), 100% laser power was used, with five iterations every five scans. The time series was designed without any intervals, with the first two time points recording the signal before bleaching, followed by 98 time points after photoconversion. Movies were processed using Airyscan and analyzed for various parameters.

### Image analyses

All Z-stacked images or movies were 3D Airyscan-processed, followed by maximum intensity projection, and saved in a czi format using Zeiss ZEN Blue software version 3.7. These projected images or movies were analyzed using FIJI or Arivis Vision4D (version 4.1). Arivis pipelines were designed as described earlier^65^. Briefly, in all images, cell bodies and non-neuronal cells were manually marked and masked out during quantification. Further, neurons were manually marked before starting the pipeline to segment objects only within the neurons.

### Zebrafish Husbandry

All zebrafish (*Danio rerio*) work was done in accordance with the University of Wisconsin-Madison Institutional Animal Care and Use Committee guidelines. Adult zebrafish were maintained at 28 °C, and embryos were spawned according to established protocols^66^. Strains used include AB (wild type; ZDB-GENO-960809-7). Embryos/larvae were kept in embryo media (995 μM MgSO_4_, 154 μM KH_2_PO_4_, 42 μM Na_2_HPO_4_, pH 7.2, 1.3 mM CaCl_2_, 503 μM KCl, 15 mM NaCl, 714 μM NaHCO_3_), maintained at 28 °C, and developmentally staged using established methods^67^. All experiments were performed at 4 days post-fertilization (dpf) unless otherwise specified. Sex is not determined at this stage.

### Transient transgenesis and live imaging

Transient transgenesis was used to express all transgenes for analysis, as previously described^35,36^. For this, 3-13 pg of DNA was microinjected into zebrafish zygotes, and animals were raised to 4 dpf. Expression was driven by a 5-kilobase portion of the neurod promoter, as previously described^68^. Plasmids used include: 5kbneurod: Polg2-GFPp2aMitoTagRFP. After microinjection, larvae were screened at 4 dpf for posterior lateral line (pLL) neuron expression on a Zeiss AxioZoom V1.6 before live imaging. For all live imaging, larvae were anesthetized in 0.02% tricaine, mounted in 1.5% low-melt agarose in embryo media, and imaged with an Olympus FV3000 confocal microscope with a 40x (NA 1.25) silicone oil objective. An optimal interslice interval was used for all imaging.

Gibson cloning was used to generate expression constructs. Expression was limited to the mitochondrial matrix (Mito) using a mitochondrial localization sequence derived from Cox8a^69^. The human Polg2-GFP ortholog has been described previously^8^. These transgenes were subcloned using PCR-based Gibson cloning to generate constructs for expression in zebrafish neurons. Simultaneous expression was promoted using the cleavable p2a peptide linker. All plasmids were sequence verified before use.

### Zebrafish POLG2-GFPanalysis

Polg2 punctal volume relative to mitochondrial volume and mitochondrial ATPSnFR signal were quantified in single pLL axon branchpoints and the axon proximal axon. Branchpoints were defined as the region of the pLL axon between the pLL nerve and where the axon then splits into secondary branches to innervate sensory cells. The region of the same axon in the pLL nerve proximal to the branch point was analyzed for comparison. Z-stacks were taken through these regions as described above with optimal imaging parameters to maximize signal-to-noise. For Polg2 analysis, the ImageJ Image Calculator plugin was used to isolate mitochondrial Polg2 and quantify the volume of Polg2 relative to mitochondrial volume. For this, the mitochondrial TagRFP was used as a mask first to isolate and then quantify the mitochondrial Polg2 punctal volume using the 3D Objects Counter plugin. Mitochondrial volume was then measured similarly.

### Statistical analyses related to zebrafish imaging

All image analysis was performed using ImageJ. Statistical analysis was performed using JMP 18, and data plots were created using GraphPad Prism 10. Sample size estimated based on previous work and determined by power analyses. All experiments were replicated at least 2 times (experimental replicates), with sample sizes for each experiment representing biological replicates. ANOVA and Wilcoxon/Kruskal-Wallis analysis were used for parametric and nonparametric tests, respectively.

### Statistical analyses related to mouse imaging

Detailed descriptions of the sample size, statistical test, and p-values are mentioned in the figure legends. All the statistical analyses were performed using GraphPad Prism. For comparison between two samples, a t-test with a 95% confidence interval was used. For multiple samples, one-way ANOVA analysis was performed.

## Data availability

All data needed to evaluate the conclusions in the paper are present in the paper and/or the Extended Data.

## Acknowledgments

The authors thank Uyan Vo for training in mouse DRG dissection.

## Funding

National Institutes of Health grant R35GM147218 (S.C.L.)

National Institutes of Health grant R01NS124692 (C.D.)

National Institutes of Health Training Grant T32GM148378 (C.Z.)

National Science Foundation grant MCB-2339182 (S.C.L.)

National Science Foundation grant DEG-2137424 (A.E.L.)

UC Berkeley Chancellor’s Predoctoral Fellowship (C.Z.)

Sloan Research Fellowship in Neuroscience (S.C.L.)

United Mitochondrial Disease Foundation Postdoctoral Fellowship (T.P.W.)

Rennie Fund for the Study of Epilepsy (S.C.L.)

Howard Hughes Medical Institute Investigator (D. B.)

## Author information and contribution

Conceptualization: T.P.W., S.C.L.

Data acquisition and analysis: T.P.W., S.C.L., A.E.L., C.D., C. V., A.S., C.Z.

Funding acquisition: T.P.W., S.C.L., A.E.L., C.D., D. B., C.Z.

Resources: S.C.L., C.D., D. B., C. V., A.S.

Writing: T.P.W., S.C.L.

Administration: C. V., A.S.

