## Extended Data Figures 1-5 for "Self-renewal of neuronal mitochondria through asymmetric division"

Extended Data Fig.1

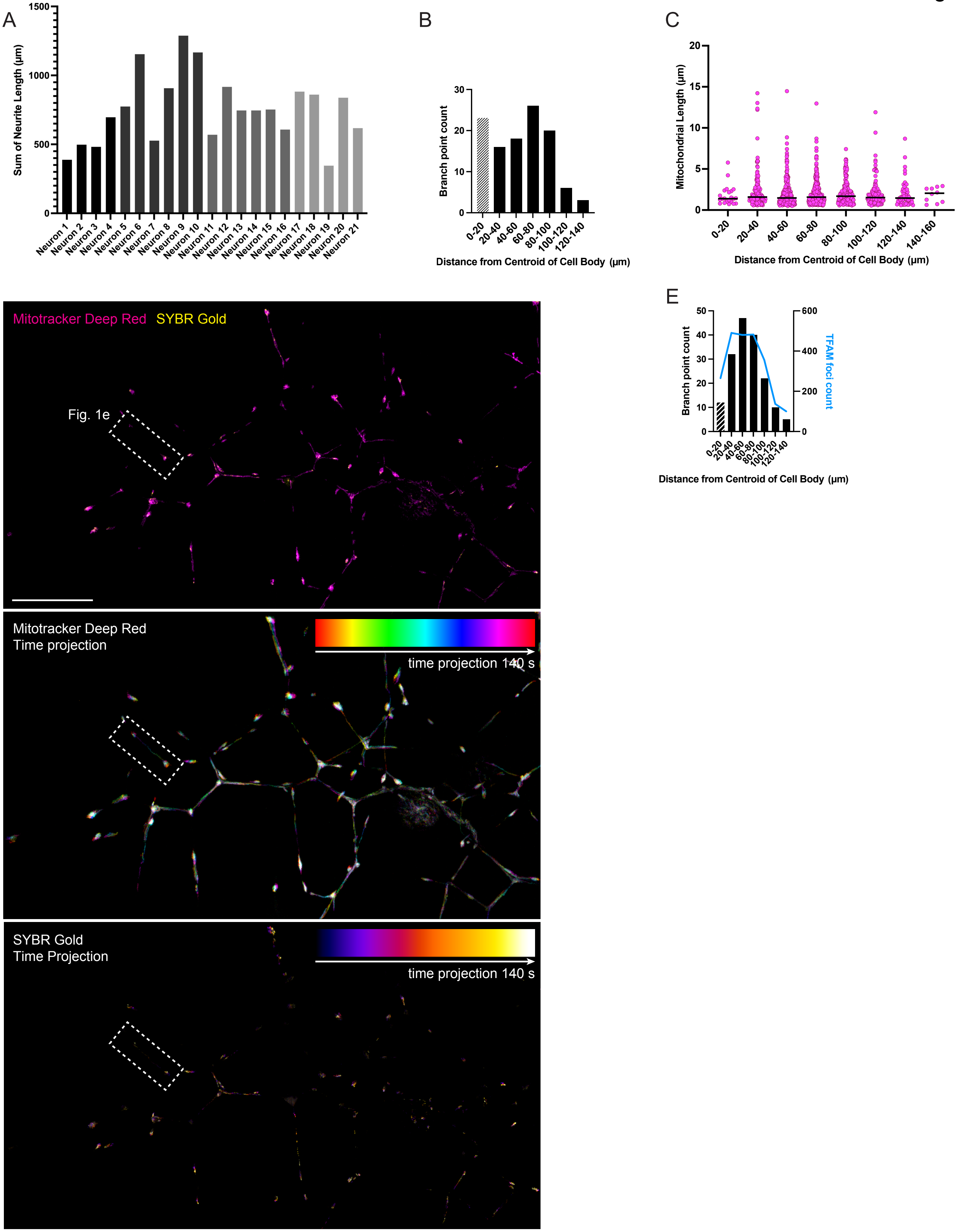

Extended Data Fig.2

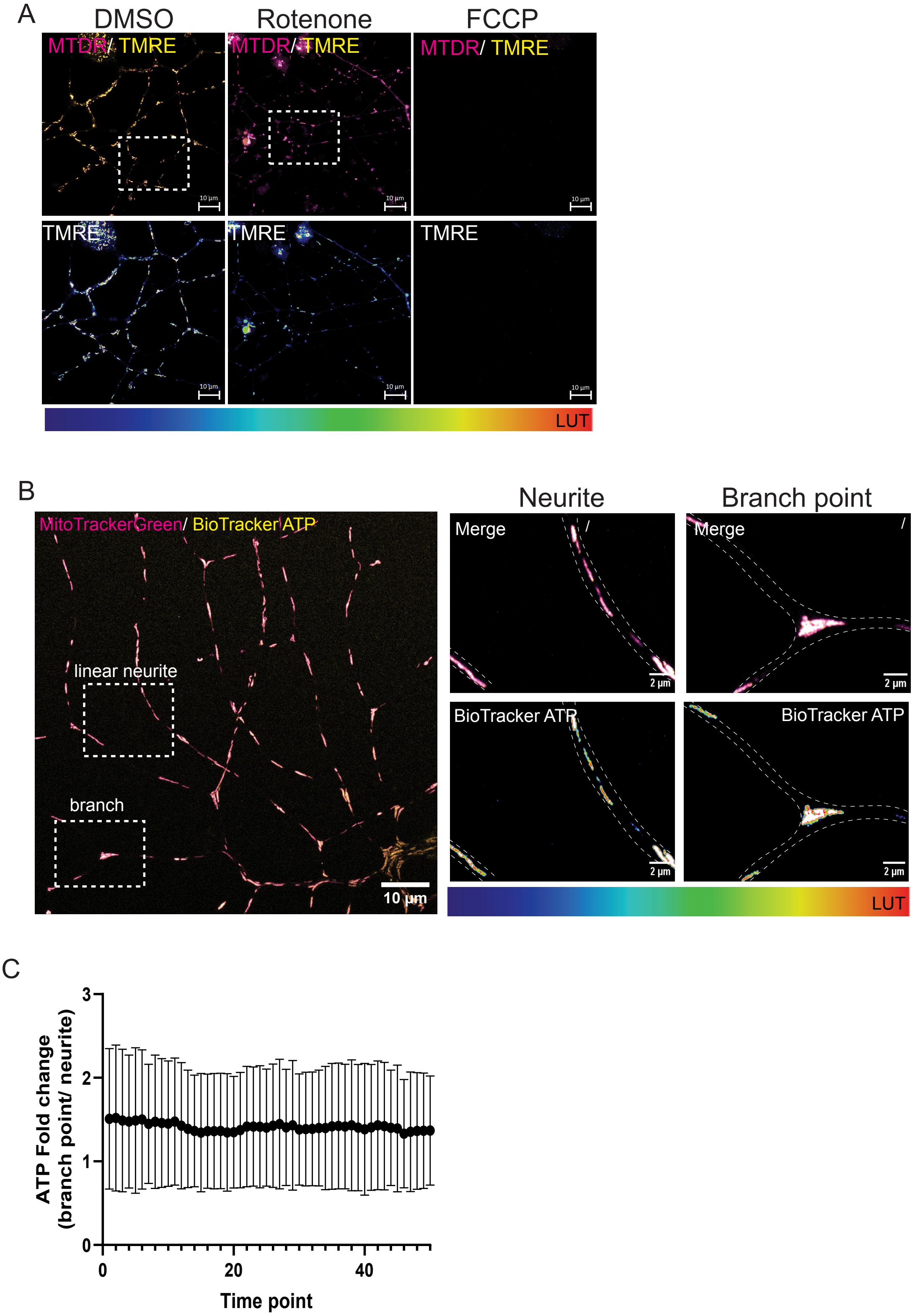

Extended Data Fig.3

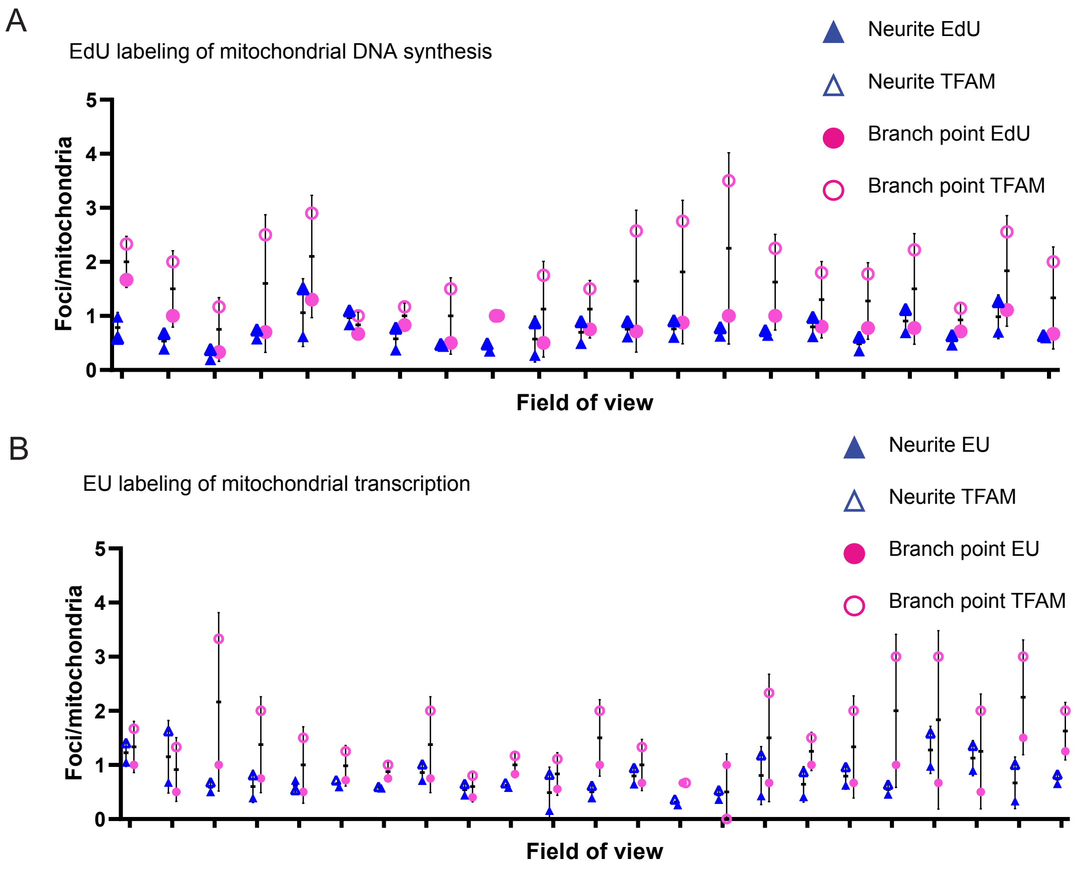

### Extended Data Fig.4

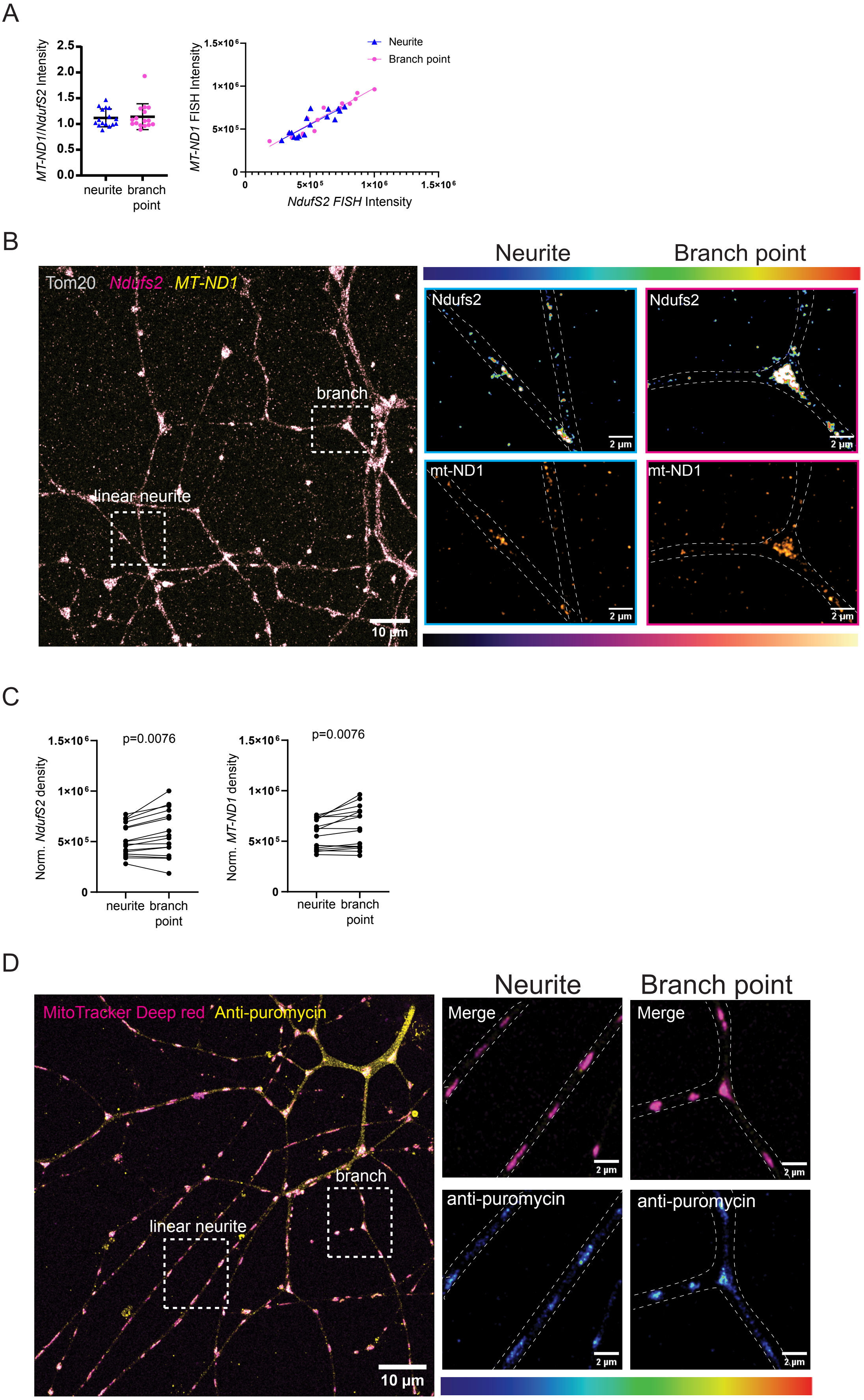

### Extended Data Fig.5

A

Classes of mitochondrial dynamic events observed in neurites

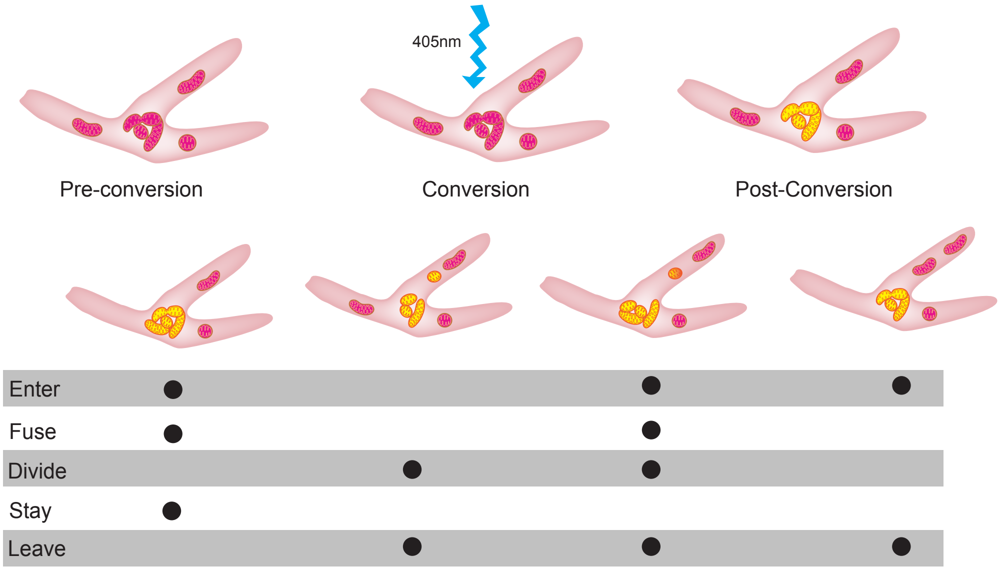


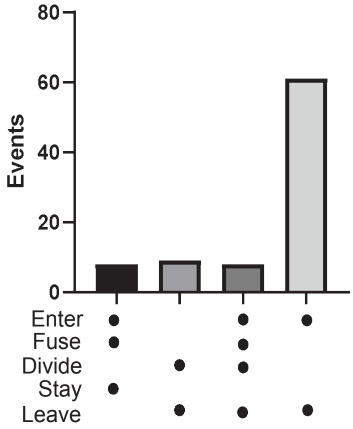


B

Mitochondrial division frequency and mtDNA partitioning

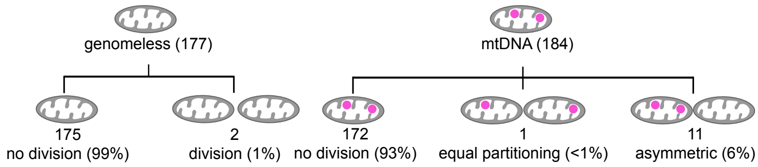
