## Supplementary material for "Self-renewal of neuronal mitochondria through asymmetric division": Table S5

| Name | Sequence |
|---|---|
| Mouse mt-ND1_1 | acgaggagtgttaggatatt |
| Mouse mt-ND1_2 | ggctatggcgattagaatgg |
| Mouse mt-ND1_3 | tgcgttctactaatgttagg |
| Mouse mt-ND1_4 | agttgtatgtaccctaagat |
| Mouse mt-ND1_5 | caacaatgttagggcctttt |
| Mouse mt-ND1_6 | ggttgtaaaatgccgtatgg |
| Mouse mt-ND1_7 | ataattttatggcgtctgca |
| Mouse mt-ND1_8 | aaagggcgtattggttcttt |
| Mouse mt-ND1_9 | agtgatagggtaggtgcaat |
| Mouse mt-ND1_10 | ctcatagacttaatgctagt |
| Mouse mt-ND1_11 | ttaatgggtgtggtattggt |
| Mouse mt-ND1_12 | actgataggctagatgttgc |
| Mouse mt-ND1_13 | tcctgatcatagaatggagt |
| Mouse mt-ND1_14 | agtatttggagtttgaggct |
| Mouse mt-ND1_15 | gctcgtaaagctccgaatag |
| Mouse mt-ND1_16 | gctatggttacttcatatga |
| Mouse mt-ND1_17 | gtagagagtaggatccattt |
| Mouse mt-ND1_18 | gttcttgggttgtaataagt |
| Mouse mt-ND1_19 | ggctggcagaagtaatcata |
| Mouse mt-ND1_20 | atcatattatggctatgggt |
| Mouse mt-ND1_21 | ttgtttctgctagggttgag |
| Mouse mt-ND1_22 | aattctgattctccttctgt |
| Mouse mt-ND1_23 | gcgtattctacgttaaaccc |
| Mouse mt-ND1_24 | agaataacgcgaatgggccg |
| Mouse mt-ND1_25 | atgttagtgtactctgctat |
| Mouse mt-ND1_26 | gttgttagggcgtttattag |
| Mouse mt-ND1_27 | tatagtataggggtcctagg |
| Mouse mt-ND1_28 | agttgagtagagttctggta |
| Mouse mt-ND1_29 | atgatagtagtagagcttct |
| Mouse mt-ND1_30 | atgctcggatccataggaat |
| Mouse mt-ND1_31 | atcgtaacggaagcgtggat |
| Mouse mt-ND1_32 | aggggtagaaagttttttca |
| Mouse mt-ND1_33 | gtcacatacataatgctagt |
| Mouse mt-ND1_34 | ggtactcccgctgtaaaaat |
| mouse Ndufs2_1 | attcaatatctggctgccac |
| mouse Ndufs2_2 | agctcccgaaaactgctctg |
| mouse Ndufs2_3 | tttccttggaggggtacatg |
| mouse Ndufs2_4 | cacatcattccaaggaggag |
| mouse Ndufs2_5 | caccgctttttccttcaaaa |
| mouse Ndufs2_6 | ccaaagttcagggtcatgtt |
| mouse Ndufs2_7 | cagcacgagtctcaggactc |
| mouse Ndufs2_8 | cgatgtgagggtcacatttc |
| mouse Ndufs2_9 | tgtactcaatgagcttctcc |
| mouse Ndufs2_10 | atggaagggcctgcagatag |
| mouse Ndufs2_11 | acatagtccaaccggtcaaa |
| mouse Ndufs2_12 | cctgttcattacacatcatg |
| mouse Ndufs2_13 | ttctccacagctatcgaata |
| mouse Ndufs2_14 | ggaggaggttggatgtttag |
| mouse Ndufs2_15 | aaagagcactcggatccact |

| | |
|---|---|
| mouse Ndufs2_16 | ttaaaatccgtgtgatctct |
| mouse Ndufs2_17 | tgtggtgacagccatgatat |
| mouse Ndufs2_18 | aagaaaggagtcatggcacc |
| mouse Ndufs2_19 | acccgctcatagaactcgaa |
| mouse Ndufs2_20 | aatgtcatccagaagcccaa |
| mouse Ndufs2_21 | cacctcatcaatccgaagag |
| mouse Ndufs2_22 | gattctattgttggtcagca |
| mouse Ndufs2_23 | caatgtcgactgtcctattt |
| mouse Ndufs2_24 | cccactgaatccatagttaa |
| mouse Ndufs2_25 | cctggtcgtaaacgtcatag |
| mouse Ndufs2_26 | cctcgagaaccgataggaac |
| mouse Ndufs2_27 | gacacaggtacctatcgtag |
| mouse Ndufs2_28 | tcaatgattcgcagggactg |
| mouse Ndufs2_29 | gaggcatcttgttcagacac |
| mouse Ndufs2_30 | gacactttggcgtcatcaac |
| mouse Ndufs2_31 | gacgtcttcatctctgctcg |
| mouse Ndufs2_32 | gtgatgaattagtgactcca |
| mouse Ndufs2_33 | aacttggtagccctctgtat |
| mouse Ndufs2_34 | aatggcagtatatgtggctc |
| mouse Ndufs2_35 | caaactctcccttaggagct |
| mouse Ndufs2_36 | ccatcggataccaagtatac |
| mouse Ndufs2_37 | gatcttacaccgataagggc |
| mouse Ndufs2_38 | caggtgggcaaaaccgggag |
| mouse Ndufs2_39 | ccttagacatcttgtccaaa |
| mouse Ndufs2_40 | acgacatctgccaacatgtg |
| mouse Ndufs2_41 | aatatcctgggtacctatga |
| mouse Ndufs2_42 | atcggtctatttctccgaac |
| mouse Ndufs2_43 | ctcgaaggagcagctgacag |
| mouse Ndufs2_44 | agaaagcctgcgcagctaag |
| mouse Ndufs2_45 | tgtctccttttctagttcat |
| mouse Ndufs2_46 | ggccaaaaggtggctaattt |
